# The genetic basis of traits associated with the evolution of serpentine endemism in monkeyflowers

**DOI:** 10.1101/2022.12.11.519970

**Authors:** Katherine Toll, John H. Willis

## Abstract

The floras on chemically and physically challenging soils, such as gypsum, shale, and serpentine, are characterized by narrowly endemic species. These edaphic endemics often have widespread close relatives that are not restricted to specific soil types. The evolution of edaphic endemics may be facilitated or constrained by genetic correlations among traits contributing to adaptation and reproductive isolation across soil boundaries. The yellow monkeyflowers in the *Mimulus guttatus* species complex are an ideal system in which to examine these evolutionary patterns. To determine the genetic basis of adaptive and prezygotic isolating traits, we performed genetic mapping experiments with F2 hybrids derived from a cross between a serpentine endemic, *M. nudatus*, and its close relative *M. guttatus*. Plants occurring on serpentine soils have repeatedly evolved short statures and small leaves, suggesting that these traits are adaptive, and *M. nudatus* shows all these characteristics compared to *M. guttatus*. Previous research demonstrated that flower size and life history differences between these species contribute to prezygotic reproductive isolation between them. Few large effect and many small effect loci contribute to interspecific divergence in life history, floral and leaf traits, and a history of directional selection contributed to trait divergence. Loci contributing to adaptive traits and prezygotic reproductive isolation overlap, and their allelic effects are largely in the direction of species divergence. These loci contain promising candidate genes regulating flowering time and plant organ size. Together our results suggest that genetic correlations among traits facilitated the evolution of edaphic adaptation and speciation in this species pair.

## Introduction

Geology influences the composition, abundance, and traits of species, exemplified by dramatic floristic transitions across soils. Soils that are chemically and physically challenging to plant species, like shale barrens, granite outcrops and serpentine, typically harbor unique floras characterized by narrowly endemic species (Wherry, 1930; McVaugh, 1943; Whittaker, 1954). These harsh soils can facilitate the persistence of older species as refugia and the evolution of new species (Stebbins, 1942; Stebbins & Major, 1965). Many edaphic (soil-related) endemics have close relatives that occur nearby and are not restricted to specific soil types, suggesting that new species often evolve from widespread progenitors across soil boundaries (Anacker et al., 2011; Grossenbacher et al., 2014; Anacker & Strauss, 2014). Yet edaphic endemics are typically outnumbered by widespread species on their specialized soils, which may reflect decreased diversification rates in endemics relative to widespread species (Anacker et al., 2011) or the difficulty of evolving reproductive isolation with gene flow (Caisse & Antonovics, 1978; Felsenstein, 1981). The genetic architecture of divergence may influence the probability of speciation across soil boundaries and coexistence upon secondary contact. If adaptive and reproductive isolating traits have a shared genetic basis, genetic correlations, either through pleiotropy or linkage, can facilitate the formation and maintenance of new species across soil boundaries (Hawthorne and Via, 2001). However, few studies have investigated the number, location, and effect size of loci contributing to reproductive isolation and adaptive divergence between edaphic endemics and their close relatives (Macnair & Cumbes, 1989; Ferris et al., 2017).

Whether endemics arise through divergence of individuals that establish in a new edaphic environment from a widespread ancestral species or depletion of a previously widespread species composed of edaphic ecotypes, both routes to endemism begin with the evolution of locally adapted populations. Local adaptation is common in plant populations inhabiting different edaphic environments (Wright et al., 2006; Sambatti & Rice, 2006; Hereford, 2009) and can evolve rapidly (Wu et al., 1975; Davies & Snaydon, 1976). Classic evolutionary theory predicts that most adaptive substitutions have small or intermediate effects on fitness due to pleiotropy (Fisher, 1930) and drift (Kimura, 1983). More recent theory predicts an exponentially decreasing distribution of effect sizes when populations are far from their phenotypic optimum (Orr, 1998a), and that large effect loci play a major role in adaptation in the presence of gene flow (Griswold, 2006; Yeaman & Whitlock, 2011; reviewed in Dittmar et al., 2016), conditions that are likely in the evolution of local adaptation across soil boundaries. In addition, soil boundaries are often discrete, transitioning abruptly between soil types with no intermediates, a scenario where loci with small effects may be selected against because intermediate phenotypes have low fitness (MacNair, 1983; Selby & Willis, 2018). While adaptation across soil boundaries may initially favor the fixation of alleles with large effects (e.g., loci contributing to soil tolerance), adaptive alleles subsequently fixed within edaphic endemics may have smaller effects for several reasons. First, as locally adapted populations evolve reproductive isolation, gene flow becomes less likely to swamp divergence at smaller effect alleles. Furthermore, environmental differences may no longer be as extreme and abrupt as between soil types, and thus smaller effect alleles may be favored.

Edaphic ecotypes often differ in traits contributing to prezygotic reproductive isolation, like flowering time and mating system (McNeilly & Antonovics, 1968; Dittmar & Schemske, 2018; Sianta & Kay, 2021). Genetic correlations among adaptive and reproductive isolating traits may facilitate divergence between soil ecotypes and may be common between taxa inhabiting chemically and physically different soils. For example, loci contributing to serpentine soil adaptation (nickel tolerance, leaf area and succulence) and premating isolation (flowering time) overlap between *Silene vulgaris* ecotypes (Bratteler et al., 2006). Loci contributing to divergence in adaptive (leaf shape) and prezygotic isolating traits (flowering time and flower size) overlap between a granite endemic and its widespread close relative (Ferris et al., 2017).

The *Mimulus guttatus* [syn. *Erthranthe guttata* (Fisch. DC.) G.L. Nesom] species complex, a group of closely related yet morphologically and ecologically diverse wildflowers, is an excellent system for investigating the genetic basis of adaptation and reproductive isolation across soil boundaries. *M. guttatus* is widespread throughout western North America and is closely related to multiple recently derived geographically restricted species nested within its range. These close relatives are often edaphic endemics, including on granite, serpentine, and copper mines, suggesting that speciation occurs repeatedly across soil boundaries in this group (MacNair et al., 1989; Ferris et al., 2014; Selby et al., 2014). One example is *M. nudatus*, an annual mixed-mating serpentine soil endemic restricted to the northern coast range of California. Serpentine soils, derived from metamorphic ultramafic rock, have low levels of Ca relative to Mg, are deficient in essential plant nutrients, often have an excess of toxic heavy metals, are rocky, shallow, and have a low water holding capacity (Brady et al., 2005). Populations of *M. guttatus* have repeatedly evolved serpentine tolerance (Selby & Willis, 2018), and these annual mixed-mating serpentine tolerant populations co-occur within meters of populations of *M. nudatus* (Gardner & MacNair, 2000; Toll & Willis, 2018). Throughout its range, *M. guttatus* lives in streams and meadows, whereas *M. nudatus* grows in gravel washes and rocky outcrops. Consistent with their habitat affinities, in lab experiments *M. nudatus* is more drought tolerant than *M. guttatus* (Hughes et al., 2001; Wu et al., 2010). The drier sites inhabited by *M. nudatus* tend to have fewer competitors and more bare ground than the more densely vegetated and wetter sites inhabited by *M. guttatus*, a common pattern observed in serpentine soil endemics when compared to species that live on and off serpentine soils (Sianta & Kay, 2019).

Plants adapted to serpentine soils usually have smaller leaves and shorter statures compared to non-serpentine populations or closely related species (Pichi-Sermolli, 1948; Rune, 1953; reviewed in Krukeberg, 1954). While serpentine adapted populations of *M. guttatus* are slightly shorter and have smaller rosettes than non-serpentine populations off serpentine soils, they are phenotypically very similar to off-serpentine populations (Selby & Willis, 2018). Consistent with broader floristic patterns, the serpentine endemic *M. nudatus* is smaller than both non-serpentine and serpentine-adapted populations of *M. guttatus* and differs in several ecologically important traits: *M. nudatus* flowers under shorter critical daylengths and often flowers earlier in the greenhouse, has smaller flowers, and smaller and narrower leaves than *M. guttatus* (Figure 1, Selby et al., 2014; Freidman & Willis, 2013; Ferris et al., 2015, Sianta & Kay, 2021). Small leaves and small flowers lose water to transpiration at a slower rate than large leaves and consequently are associated with arid environments (Galen et al., 1999; Nicotra et al., 2011), and early flowering is a common drought escape strategy in plants (Kooyers, 2015). Early flowering in *M. nudatus* is associated with an earlier developmental shift in allocation from vegetative to reproductive growth – *M. nudatus* transitions to reproductive growth and flowers at an earlier developmental stage, reflected by earlier bolting (elongation between earlier forming leaf pairs) and flowering at earlier nodes (leaf pairs), respectively. Finally, while hybrid seed inviability is an extremely important and strong postzygotic barrier between these species (Gardner & McNair 2000; Oneal et al., 2016; Toll & Willis, 2018), other reproductive barriers maintain species boundaries and facilitate coexistence in this species pair. Specifically, flowering time and flower size differences contribute to temporal and mechanical prezygotic isolation between them in sympatry (Gardner & MacNair, 2000; Sianta & Kay, 2021).

**Figure 1.**
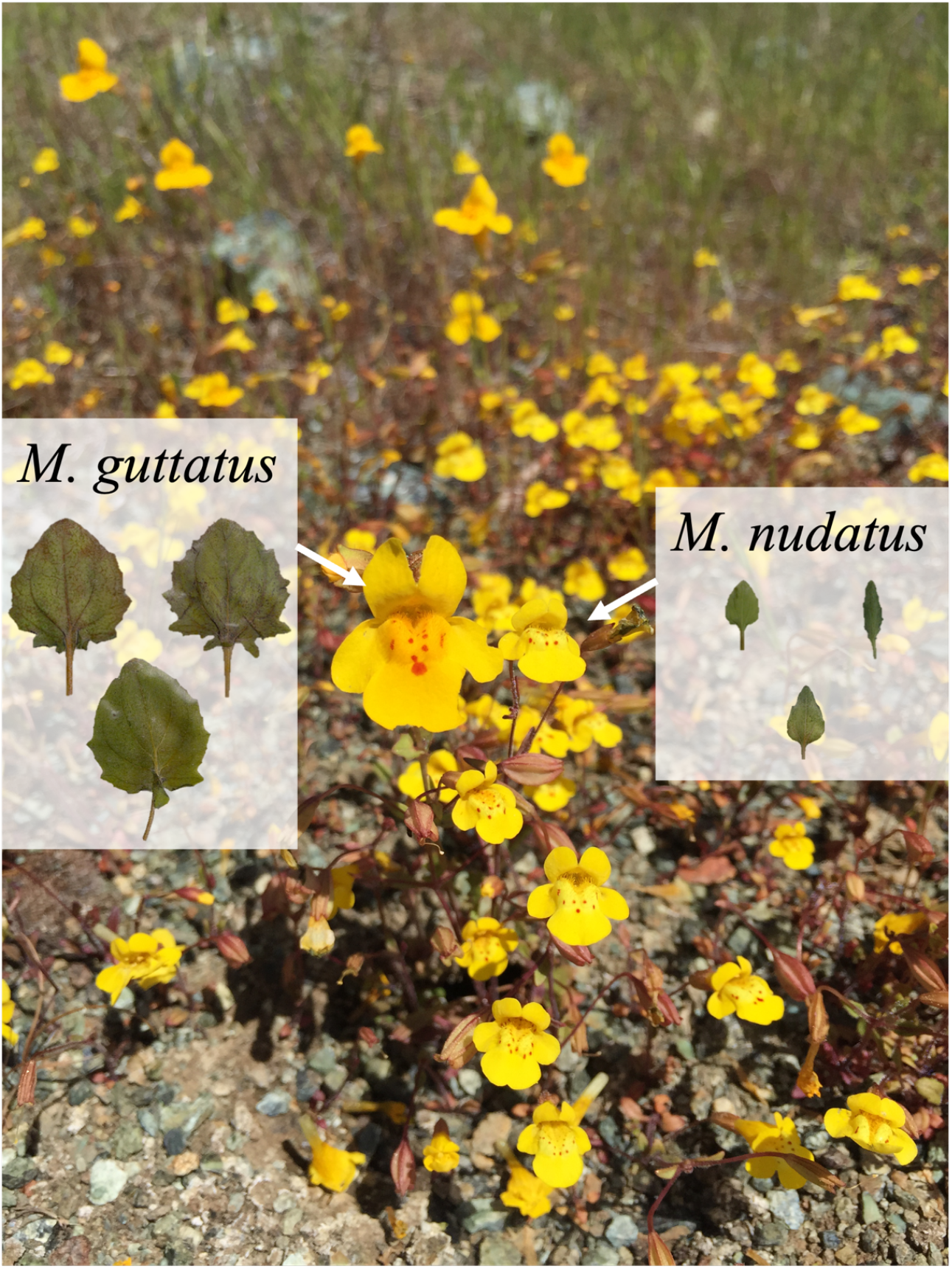
Photograph of co-occurring *Mimulus guttatus* and *Mimulus nudatus*. Leaves were scanned from individuals belonging to 3 co-occurring populations for each species.

To determine the genetic architecture of traits contributing to adaptive divergence and prezygotic reproductive isolation between *M. nudatus* and *M. guttatus*, we performed a genetic mapping experiment with F2 hybrids derived from a cross between them. We crossed lines from a sympatric serpentine tolerant population of *M. guttatus* to *M. nudatus*. Since serpentine and non-serpentine populations of *M. guttatus* are phenotypically very similar, this allowed us to focus on traits associated with occupation of drier and sparser microhabitats typical of serpentine endemics. Furthermore, prezygotic isolating traits between these species are more important in sympatry, where soil boundaries do not restrict hybridization. With this experiment we asked the following questions: What is the genetic architecture of prezygotic reproductive isolation and putatively adaptive trait divergence associated with serpentine endemism? Do the loci contributing to reproductive isolation and adaptive traits overlap? Are allelic effects at overlapping loci all in the direction of species divergence (facilitating divergence) or do they oppose each other (constraining divergence)? Did a history of consistent directional selection contribute to species divergence?

## Materials and methods

### Plant material and experimental design

We created inbred lines from maternal families collected at a sympatric site at the UC Donald and Sylvia McLaughlin Natural Reserve in Lower Lake, California (W 122° 24.614, N 38° 51.528). We created F1 hybrids from parental lines that were self-fertilized for two generations in the greenhouse, and then selfed single F1 hybrid to generate F2 hybrids. We planted seeds of the *M. guttatus* line REM122 (*n* = 21), the *M. nudatus* parental line REMn129 (*n* = 24), F1 hybrids (*n* = 10), and F2 hybrids (*n* = 576) in 2.5” Kord pots with Fafard 4P soil, cold stratified them for 5 days, and placed in a 16 hr light/8 hr dark greenhouse at Duke University. We fully randomized pots and flats were rotated weekly. Of the seeds we planted, 18 *M. guttatus*, 16 *M. nudatus*, 3 F1s and 398 F2 hybrids germinated and were included in our phenotype analysis. Of these F2s, we extracted sufficient high-quality DNA from 379 F2 hybrids to generate reduced representation libraries for QTL mapping.

### Phenotypic measurements and analyses

We recorded germination date and flowering date, and calculated days to flowering by subtracting germination date from flowering date. On the date of first flower, we recorded the node of first flower, and measured the length of the first internode, plant height, corolla width, corolla length with a metal ruler in 1/100ths of an inch. The node of first flower is the leaf pair at which a given plant produced its first flower, counting from the first true leaf pair after the cotyledons (0=cotyledon node, 1 = first true leaf node, and so on). The length of the first internode is the length between the cotyledons and the first leaf pair and describes whether a plant bolted (transitioned to flowering) early in development or not. We transformed these measurements to centimeters in our analysis. We removed the first leaf from each plant on the date of first flower, and measured leaf traits in centimeters (length, width, and area) with images produced using a flatbed scanner in ImageJ (Schneider et al., 2012). We calculated leaf shape by dividing leaf width from leaf length.

We tested whether the parental lines differed in the traits using Welch’s two sample *t*-tests, and calculated Bonferroni adjusted *p*-values by dividing each *p*-value by the number of *t*-tests (8). To estimate trait heritabilities, we calculated parental and F2s variances to estimate environmental, phenotypic, and genetic variances. First, we calculated the environmental variance as the weighted average of the parental line variances, excluding F1s due to low sample size (F1 *n*=3) (*V*_E_= (var(REM122)+var(REMn129))/2). Then, we calculated genetic variance by subtracting *V*_E_ from *V*_P_, the F2 phenotypic variance (*V*_G_= *V*_P_ – *V*_E_; Falconer & MacKay, 1996). Finally, we estimated broad-sense heritability by dividing the genetic variance by the F2 variance (*H^2^* = V_G_/V_P_). We estimated trait correlations in the F2 hybrids by calculating Pearson correlation coefficients.

### DNA extractions and library generation

We extracted DNA from F2 hybrids and parents and prepared reduced representation libraries for sequencing. Flower bud tissue was collected from each parent and F2 hybrid in 96-well plates and frozen at −80°C. DNA was extracted using a modified CTAB protocol (Lin & Ritland, 1995). Prior to grinding in a Geno/Grinder 2000, a single stainless steel grinding ball was added to each sample and flash frozen with liquid nitrogen. We modified Lin & Ritland (1995) by adding 0.28mg of RNaseA to each sample after the chloroform-isoamyl extraction, followed by water bath incubation for 30 min at 37°C, and an additional chloroform-isoamyl extraction to remove RNaseA. After the second chloroform-isoamyl extraction, we precipitated DNA using 5M NaCl and 95% ethanol. Pellets were cleaned using 70% ethanol, and then DNA pellets were dried overnight and re-suspended in autoclaved distilled water.

We determined the concentration of DNA in each sample using the Quant-iT PicoGreen dsDNA Assay kit (Invitrogen Life Technologies) and assayed fluorescence on a microplate reader. After quantification, we diluted 379 F2 samples to a standard concentration of 5ng DNA per μL of water. We then prepared the standardized samples for Illumina sequencing using a modified multiplexed shotgun genotyping (MSG) protocol (Andolfatto et al., 2011). Briefly, we digested 50ng of genomic DNA for each F2 with the restriction enzyme Csp6I and ligated 48 unique barcodes to each sample. We pooled the barcoded DNA into 8 groups of up to 48 barcoded individuals and size selected the pools using Ampure XP beads and a gel. We then amplified the pooled DNA and ligated 8 unique Illumina index primers with 16 cycles of polymerase chain reaction (PCR). Finally, we pooled equal molar amounts of the 8 Illumina indexed libraries based on concentrations measured using the Qubit broad range assay and insert size distributions measured using Agilent High Sensitivity D1000 ScreenTape. The pooled F2 hybrids were then sequenced on a single lane of Illumina HiSeq 2000/2500 with 50bp single end reads.

Parental samples were prepared by digesting 200ng of DNA with Csp6I, followed by barcoding. Libraries were size selected with Ampure XP beads using a 0.8:1 bead to sample ratio to remove fragments smaller than 300bp prior to and following library amplification. Parental lines were sequenced with four other samples on an Illumina Hi-Seq 4000 with 50bp single end reads.

### Marker generation

To create markers for genetic mapping from our sequencing data, we used the TASSEL 5.0 GBS v2 pipeline to parse individuals by barcode, align each tag to the *M. guttatus* v2 reference genome using bwa, and identify segregating SNPs (Li & Durbin, 2009; Hellsten et al., 2013; Glaubitz et al., 2014). We used default settings in TASSEL GBS v2, e.g., a minimum locus coverage of 0.1, except that we used a minimum kmer length of 45 and a minimum Illumina quality score of 20. Parental allele assignment and imputation was performed using the window LD algorithm in FSFhap with default settings, except that we did not use heterozygous calls (Swarts et al. 2014).

### Linkage mapping and QTL mapping

Linkage map construction was performed using the *onemap* package in R with markers generated by FSFhap in TASSEL 5.0 (Margarido et al. 2007). We first removed redundant markers by binning markers with pairwise recombination fractions of 0, reducing our marker number from 4737 to 1011 for linkage map construction and QTL mapping. We then grouped markers into linkage groups (LG) using a minimum logarithm of odds (LOD) score of 6 and maximum recombination fraction of 0.35 to declare linkage. We ordered markers along each chromosome using a two-point based algorithm, Rapid Chain Delineation (RCD) (Doerge 1996). We tested for segregation distortion (significant deviations from Mendelian genotype ratios) using a chi-square test and assessed significance using Bonferroni adjusted p-values.

We initially performed standard interval mapping using the *scanone* function in R/qtl, then used the *scantwo* function to perform a two-dimensional two-QTL scan using Haley-Knott regression (Broman et al., 2003). We calculated main effect and interaction penalties with 1000 permutations of the *scantwo* function to conduct stepwise fitting of multiple QTL models. We used the *stepwiseqtl* function with a maximum of 10 QTL using Haley-Knott regression to conduct a forward/backward search of models that optimized the penalized LOD score criterion. We determined the 1.5 LOD interval for each QTL using the *lodint* function in R/qtl.

We used the *fitqtl* function in R/qtl to determine the percent variance explained by each QTL, peak LOD scores, and additive (*a*) and dominance (d) effects for each significant QTL from final stepwise models for each trait. For traits where the parental lines significantly differed, we also estimate QTL effect sizes by calculating the proportion of the mean parental divergence explained by each locus. We divided the difference between the phenotype means of the homozygotes, by the difference between the phenotype means of the parents and multiplied by 100.

We calculated the degree of dominance for each QTL by dividing the absolute value of the dominance deviation (*d*) by the additive effect (*a*) for each QTL. We binned QTL in categories based on the degree of dominance estimate: additive (0-0.20), partially dominant (0.21-0.80), dominant (0.81-1.20), under- or over-dominant (> 1.20) (Stuber et al., 1987). For partially dominant and dominant loci, we examined which parental species was dominant.

In genomic regions where QTL overlap, we examined QTL effects to determine whether genetic correlations, either through pleiotropy or linkage constrain or facilitate species divergence. If overlapping QTL all have effects in the direction of the parental divergence, these genetic correlations may have facilitated divergence. On the other hand, if some overlapping QTL effects are in the direction of parental divergence while others are in the opposite direction, these correlations may have constrained divergence between this species pair.

### Tests of directional natural selection between species

We tested whether directional natural selection could explain species divergence using two tests: a QTL sign test of equal effects (QTLST-EE; Orr, 1998b) and the *v*-test (Fraser 2020). The QTLST-EE assumes that if a trait has a continuous history of directional selection, allelic effects at QTL at will largely be in the same direction as species divergence for a given trait (Orr, 1998b). Since at least six QTL must be detected to reject the null hypothesis of neutral divergence (and all six QTL would have to be in the direction of species divergence to reject the null hypothesis), we were only able to apply this test for plant height at flowering and leaf shape. The *v*-test assumes that if a trait has continuous history of directional selection, the variance between the parental lines is expected to be larger than the segregating variance in the F2 hybrids. To calculate our test statistic, *v*, we estimated the between parent variance using a one-way ANOVA, calculated the variance within each parental line and the F2s, and used the corrected equation 2 in Fraser (2020). We used a conservative value of c=2, indicating full additivity of QTL.

### Identifying candidate genes

We used the base pair positions of markers at the start and end of 1.5 LOD intervals for each QTL to identify annotated genes in the *Mimulus guttatus* v2 reference genome (Hellsten, et al. 2013). We then used The Arabidopsis Information Resource (TAIR) to examine GO annotations for genes in our QTL regions (Berardini et al., 2015). In QTL regions contributing to life history differences, we scanned for candidate genes with GO terms related to flowering time and growth hormones (gibberelins, auxins, and brassinosteroids). In QTL regions contributing to organ size and shape differences, we scanned for candidate genes with GO terms related to floral and leaf growth. We cross referenced putative candidate genes with mutant studies describing phenotypic effects of mutations at putative candidate genes.

## Results

### Parental species differed in traits associated with serpentine endemism and and reproductive isolation

The *M. nudatus* parent flowered at significantly earlier nodes, had longer first internodes, narrower and shorter corollas, and smaller and narrower leaves than the *M. guttatus* parent (Table 1, Figure 2). However, the parental lines did not significantly differ in days to flowering or height at flowering (Table 1, Figure 2). F1 hybrids were typically more similar phenotypically to the *M. guttatus* parent (Table 1, Figure 2). Trait heritabilities ranged from 0.23 for leaf area to 0.78 for first internode length (Table 1).

**Figure 2.**
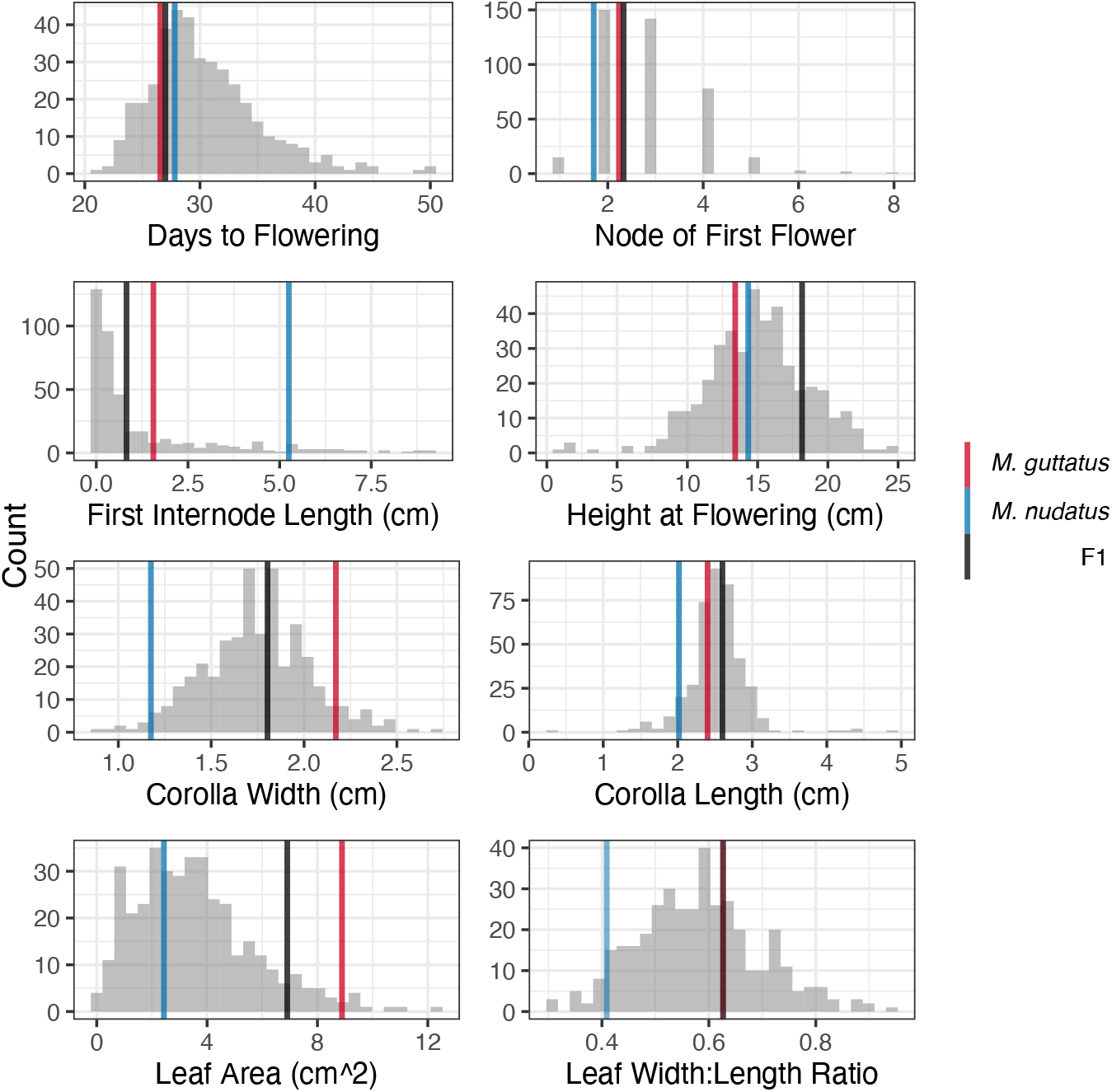
Trait distributions in the F2 mapping population between *Mimulus guttatus* and *M. nudatus*. Horizontal lines depict parental (*M. guttatus*, red; *M. nudatus*, blue) and F1 (black) means.

**Table 1.** Broad sense heritability (*H^2^)* for each trait, trait means and standard errors (SE in parentheses) for the *M. guttatus* parental line, the *M. nudatus* parental line, F1 hybrids, and F2 hybrids, *t*-test testing whether parental lines significantly differ in each trait, and Bonferonni adjusted *p*-values for the *t*-tests (significant *p*-values in bold).

| Trait | $H^2$ | *M. guttatus*<br>(n=18) | *M. nudatus*<br>(n=16) | F1 hybrids<br>(n=3) | F2 hybrids<br>(n=398) | Welch Two Sample t-<br>test comparing<br>parental lines | Bonferroni<br>adjusted p-<br>value |
| --- | --- | --- | --- | --- | --- | --- | --- |
| Days to<br>Flowering | 0.41 | 26.56<br>(1.16) | 27.81<br>(0.42) | 27.00<br>(1.15) | 30.42<br>(0.24) | $t = -1.02, df = 21.36, p$-<br>value = 0.32 | 1 |
| Node of First<br>Flower | 0.74 | 2.24 (0.14) | 1.71 (0.11) | 2.33 (0.33) | 2.88 (0.05) | $t = 2.98, df = 31.02, p$-<br>value = 0.0056 | **0.045** |
| First Internode<br>Length (cm) | 0.78 | 1.56 (0.20) | 5.25 (0.23) | 0.82 (0.55) | 1.27 (0.09) | $t = -12.15, df = 32.34, p$-<br>value = 1.34E-13 | **1.07E-12** |
| Plant Height at<br>Flowering (cm) | 0.69 | 13.41<br>(0.53) | 14.33<br>(0.46) | 18.16<br>(1.33) | 14.95<br>(0.18) | $t = -1.32, df = 31.36, p$-<br>value = 0.20 | 1 |
| Corolla<br>Width (cm) | 0.59 | 2.17 (0.05) | 1.18 (0.04) | 1.80 (0.13) | 1.74 (0.14) | $t = 15.44, df = 26.42, p$-<br>value = 9.69E-15 | **7.75E-14** |
| Corolla<br>Length (cm) | 0.49 | 2.40 (0.10) | 2.02 (0.03) | 2.60 (0.09) | 2.53 (0.02) | $t = 3.87, df = 17.16, p$-<br>value = 0.0013 | **0.010** |
| Leaf Area<br>(cm<sup>2</sup>) | 0.23 | 8.89 (0.62) | 2.44 (0.18) | 6.91 (0.41) | 3.54 (0.11) | $t = 9.95, df = 18.60, p$-<br>value = 6.99E-09 | **5.59E-08** |
| Leaf Width<br>Length Ratio | 0.73 | 0.63 (0.02) | 0.41 (0.01) | 0.63 (0.05) | 0.58 (0.01) | $t = 10.57, df = 23.57, p$-<br>value = 2.02E-10 | **1.62E-09** |

**Table 2.** QTL detected in an F2 cross between *Mimulus guttatus* and *M. nudatus:* chromosome, peak position in cM, 1.5 LOD intervals, and peak LOD score.

| Trait | Chromosome | Peak<br>Position<br>(cM) | 1.5-LOD<br>interval<br>(cM) | Peak<br>LOD score |
| --- | --- | --- | --- | --- |
| Days to<br>Flowering | 11 | 16.4 | 10.2-22.0 | 8.44 |
| Node of<br>First Flower | 10 | 41 | 34.0-56.9 | 4.17 |
|  | 11 | 23.2 | 18.0-24.9 | 19.04 |
| **First Internode Length (cm)** | 10 | 47 | 40.0-59.0 | 4.37 |
|  | 11 | 22 | 20.0-24.0 | 38.74 |
| **Height (cm)** | 2 | 37.5 | 32.1-41.0 | 5.57 |
|  | 5 | 27 | 12.0-40.0 | 6.07 |
|  | 8a | 15.8 | 0.0-23.0 | 3.94 |
|  | 8b | 70.3 | 58.1-88.0 | 3.98 |
|  | 9 | 42 | 34.0-48.0 | 7.74 |
|  | 11 | 2 | 0.0-6.0 | 4.81 |
|  | 13 | 47 | 43.2-49.0 | 15.7 |
|  | 14 | 35.1 | 33.0-39.0 | 6.52 |
| **Corolla Width (cm)** | 5 | 9 | 0.0-16.0 | 7.19 |
|  | 8a | 11.6 | 6.0-14.3 | 4.06 |
|  | 8b | 85 | 79.4-94.0 | 3.93 |
|  | 11 | 29 | 21.0-34.0 | 8.77 |
| **Corolla Length (cm)** | 11 | 27 | 21.0-32.0 | 15.06 |
| **Leaf Area (cm<sup>2</sup>)** | 5 | 7.2 | 1.1-11.0 | 5.38 |
|  | 6 | 75.6 | 71.4-77.0 | 3.95 |
| **Leaf Shape** | 4 | 64.6 | 60.0-67.0 | 8.28 |
|  | 6 | 42 | 34.0-58.0 | 5.4 |
|  | 8 | 89 | 74.0-107.2 | 4.72 |
|  | 9 | 26.8 | 13.0-32.6 | 4.89 |
|  | 12 | 44.9 | 33.0-49.0 | 4.63 |
|  | 14 | 106.7 | 98.8-108.7 | 6.22 |

### Traits associated with serpentine endemism and reproductive isolation are correlated in the F2 hybrids

Many traits measured in the F2 hybrids were significantly correlated, but correlation coefficients were not very high (Figure 3, Pearson correlation coefficients |*r*| < 0.7, per Dormann et al. 2013). The highest correlations were between node of first flower and days to flowering (*r* = 0.6), node of first flower and first internode length (*r* = −0.53), and corolla width and length (*r* = 0.53). Except for leaf shape, which was only correlated with height at flowering (*r* = −0.12), most traits were correlated with multiple (5-6) traits, suggesting a shared genetic basis among those correlated traits (Figure 3).

**Figure 3.**
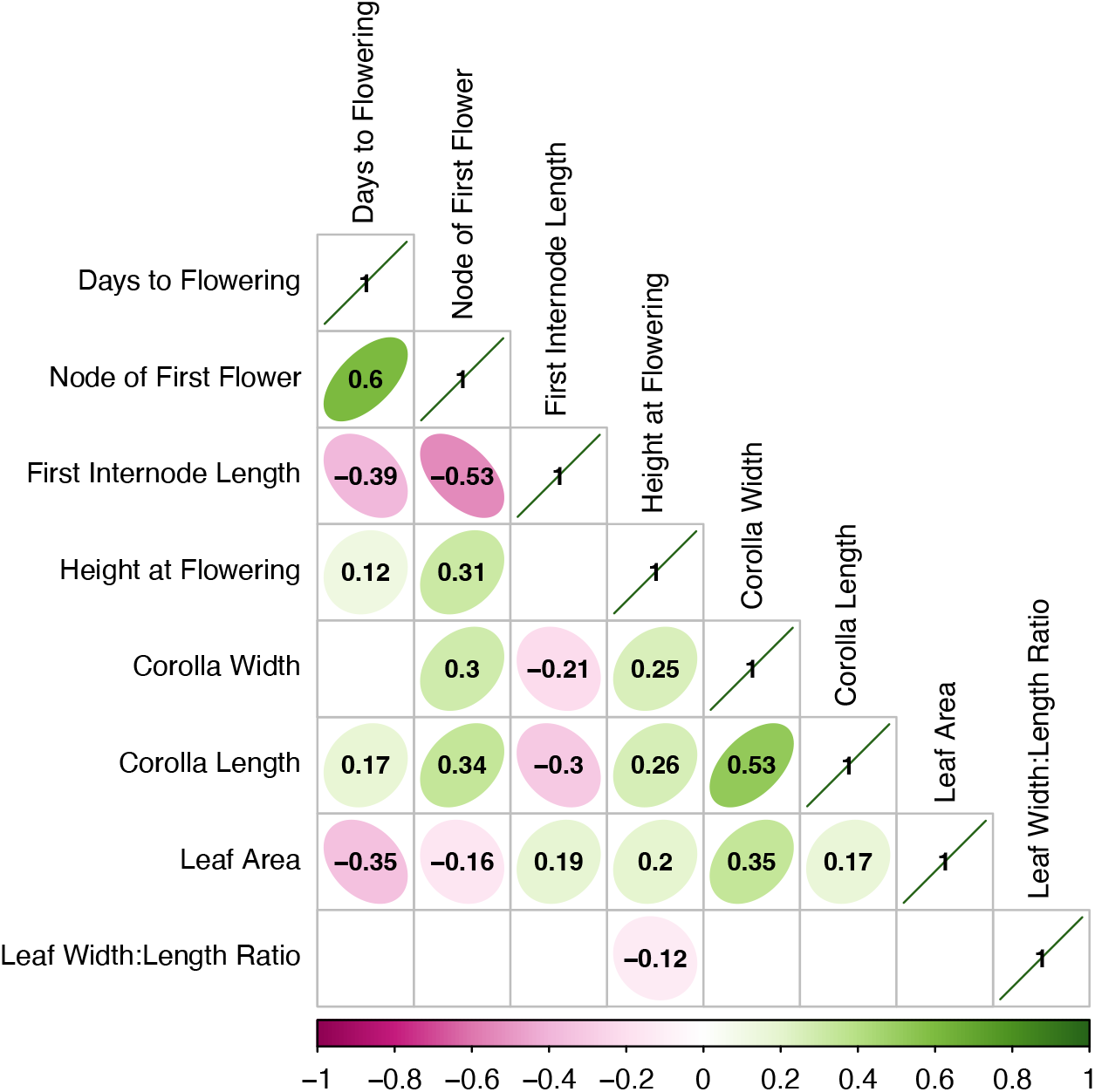
Pearson correlations among traits in the F2 hybrids. Only significant correlation coefficients are shown. The shape of each ellipse corresponds to the direction of the correlation, and color corresponds to the strength and direction of each correlation. Negative correlation coefficients are pink, and positive correlation coefficients are green.

### Loci contributing to traits associated with serpentine endemism and reproductive isolation explain a low percentage of the segregating F2 variance but a large percentage of parental divergence

Our genetic map consisted of 14 chromosomes with a total map length of 1017.5 cM (range 43.4-125.8 cM per chromosome) and average spacing of 1 cM. Roughly 22% (222/1011) of markers significantly deviated at an α of 0.05 from Mendelian ratios (1:2:1), particularly on chromosome 6 (Figure S1). Almost all distorted markers showed a deficiency of *M. nudatus* alleles relative to *M. guttatus* alleles.

We detected a total of 26 QTL across all traits, ranging from one QTL for days to flowering to eight QTL for plant height at flowering (Table 3). The average percent variance explained by an individual locus was 7.6% (range: 3% −36%), while the average total variance explained by all significant loci was 24.6% (range: 10% - 40%) for each trait (Table 3). We did not detect significant epistatic interactions for any QTL. Five QTL were additive, 14 QTL were partially dominant, three were dominant, and four QTL were over-dominant (Table 3). Of the 17 QTL that were partially dominant or dominant, 14 had an *M. guttatus* dominant allele, while only 3 had an *M. nudatus* dominant allele (Table 3).

**Table 3.** QTL positions, percent F2 trait variance explained by each locus, percent of the parental difference explained by each locus, the mean *M. guttatus* homozygote (GG) phenotype, the mean heterozygote (GN) phenotype, the mean *M. nudatus* homozygote (NN) phenotype, the additive (a) and dominance (d) effects of each locus, the degree of dominance (d/a), the mode of action (categorized based on the degree of dominance following Stuber et al. (1987), and for partially dominant and dominant loci, which allele was dominant (G= *M. guttatus*, N= *M.nudatus*).

| Trait | Chr | % Variance | % Parental Difference | Mean GG phenotype | Mean GN Phenotype | Mean NN Phenotype | a | d | Degree of dominance | Mode of action | dominant species |
| --- | --- | --- | --- | --- | --- | --- | --- | --- | --- | --- | --- |
| Days to Flowering | 11 | 9.97 |  | 32.68 | 30.74 | 27.94 | -2.36 | 0.5 | 0.21 | partial dominance | G |
| Node of First Flower | 10 | 3.98 | 113.21 | 3.08 | 2.88 | 2.48 | -0.33 | 0.17 | 0.52 | partial dominance | G |
|  | 11 | 19.93 | 267.92 | 3.51 | 2.93 | 2.09 | -0.71 | 0.08 | 0.11 | additive |  |
| First Internode Length (cm) | 10 | 3.26 | 28.18 | 1.04 | 1.1 | 2.08 | 0.52 | -0.35 | 0.67 | partial dominance | G |
|  | 11 | 35.94 | 82.38 | 0.37 | 0.8 | 3.41 | 1.6 | -1.13 | 0.71 | partial dominance | G |
| Height (cm) | 2 | 4.25 |  | 14.23 | 15.09 | 15.67 | 0.94 | 0.56 | 0.60 | partial dominance | N |
|  | 5 | 4.65 |  | 15.11 | 15.35 | 13.66 | -0.86 | 1.18 | 1.37 | overdominance |  |
|  | 8a | 2.98 |  | 13.70 | 15.00 | 16.41 | 1.00 | -0.24 | 0.24 | partial dominance | G |
|  | 8b | 3.01 |  | 13.92 | 15.12 | 15.86 | 0.83 | 0.48 | 0.58 | partial dominance | N |
|  | 9 | 5.99 |  | 16.22 | 14.90 | 14.01 | -1.35 | -0.30 | 0.22 | partial dominance | N |
|  | 11 | 3.65 |  | 14.81 | 15.59 | 13.63 | -0.36 | 1.35 | 3.75 | overdominance |  |
|  | 13 | 12.76 |  | 13.62 | 15.26 | 16.71 | 2.00 | 0.04 | 0.02 | additive |  |
|  | 14 | 5.01 |  | 15.40 | 15.10 | 13.40 | -1.26 | 0.60 | 0.48 | partial dominance | G |
| **Corolla Width (cm)** | 5 | 7.07 | 20.06 | 1.81 | 1.77 | 1.61 | -0.11 | 0.05 | 0.48 | partial dominance | G |
|  | 8a | 3.92 | 15.18 | 1.78 | 1.77 | 1.63 | -0.08 | 0.06 | 0.69 | partial dominance | G |
|  | 8b | 3.79 | 8.85 | 1.73 | 1.79 | 1.65 | -0.05 | 0.10 | 1.99 | overdominance |  |
|  | 11 | 8.71 | 21.73 | 1.81 | 1.77 | 1.59 | -0.13 | 0.12 | 0.91 | dominance | G |
| **Corolla Length (cm)** | 11 | 16.96 | 96.34 | 2.62 | 2.55 | 2.26 | -0.20 | 0.13 | 0.64 | partial dominance | G |
| **Leaf Area (cm<sup>2</sup>)** | 5 | 6.50 | 23.97 | 4.08 | 3.69 | 2.54 | -0.82 | 0.30 | 0.36 | partial dominance | G |
|  | 6 | 4.74 | -10.82 | 2.93 | 3.88 | 3.62 | 0.41 | 0.61 | 1.49 | overdominance |  |
| **Leaf Shape** | 4 | 7.91 | -39.01 | 0.54 | 0.58 | 0.62 | 0.05 | 0.00 | 0.11 | additive |  |
|  | 6 | 5.06 | 34.31 | 0.60 | 0.58 | 0.52 | -0.05 | 0.03 | 0.59 | partial dominance | G |
|  | 8 | 4.40 | 36.88 | 0.62 | 0.57 | 0.54 | -0.04 | 0.00 | 0.10 | additive |  |
|  | 9 | 4.56 | -28.44 | 0.54 | 0.58 | 0.61 | 0.04 | 0.01 | 0.15 | additive |  |
|  | 12 | 4.32 | -29.07 | 0.57 | 0.57 | 0.63 | 0.03 | -0.03 | 1.06 | dominance | G |
|  | 14 | 5.86 | 17.09 | 0.60 | 0.57 | 0.56 | 0.03 | -0.03 | 1.09 | dominance | G |

For most traits, the sum of the variance explained by all loci was less than half of the trait heritabilites (average: 41%, range: 24% - 57%), suggesting that additional loci with small effects contribute to each of our measured traits. For example, days to flowering had a broad sense heritability of 0.41 (Table 1) and a single QTL explained ~10% of the segregating F2 variance (Table 3), leaving 76% of the heritability unexplained. On the other hand, many loci individually explained a large percentage of the parental divergence, suggesting that few substitutions between species are necessary to explain dramatic phenotypic differences between them (Table 2, Figure 4). For traits where the parental lines differed, QTL effects were typically in the direction of the parental divergence (i.e., the percent of the parental divergence explained was positive for 13 out of 17 QTL in Table 3). The percent of the parental divergence explained by loci with effects in the direction of the parental divergence averaged 59% (range: 9% - 267%).

**Figure 4.**
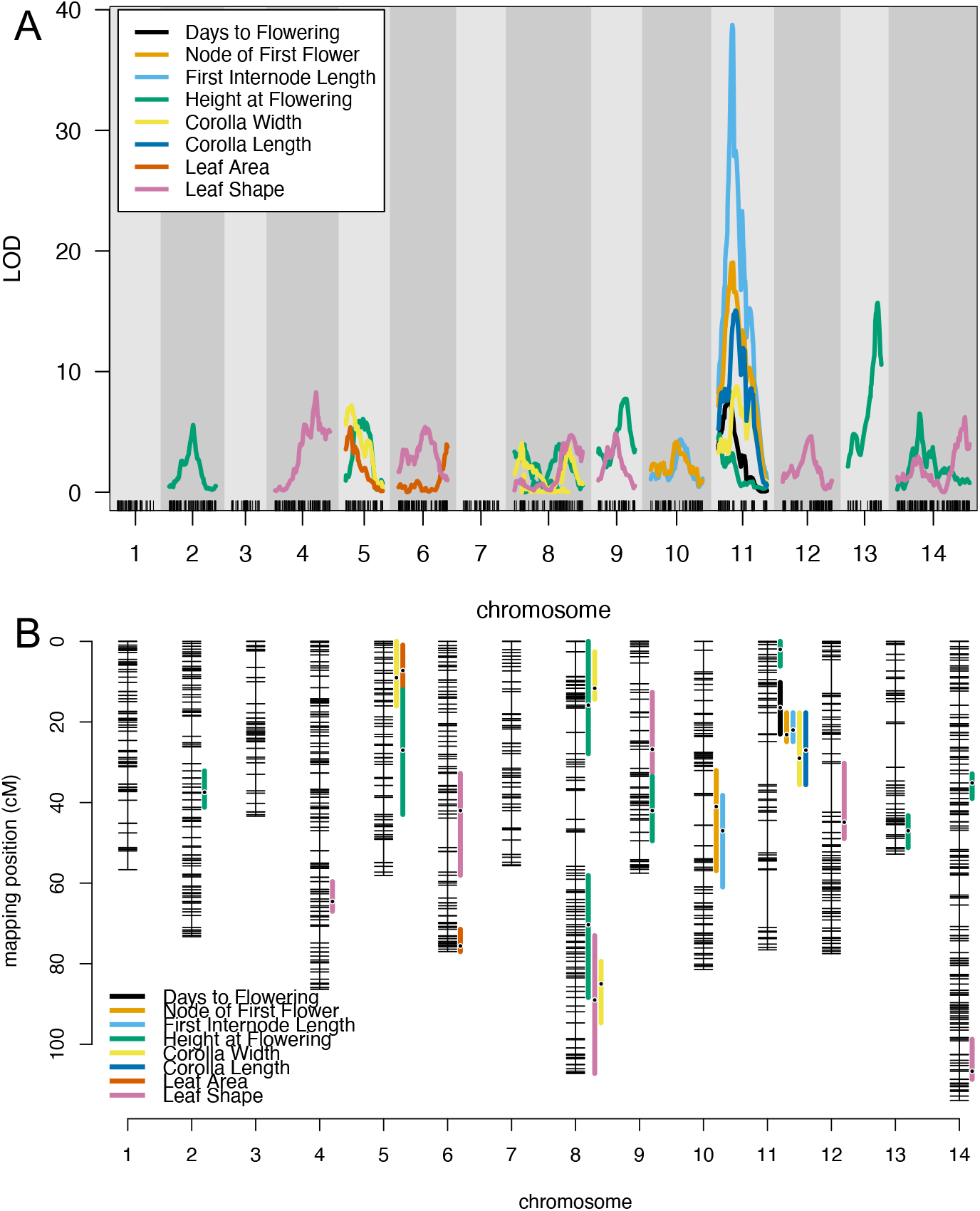
Quantitative trait loci detected across all measured traits in the F2 mapping population between *Mimulus guttatus* and *M. nudatus*. (A) Plot of significant LOD curves from multiple QTL models. (B) Plot of 1.5 LOD intervals (vertical colored bars) for each significant QTL from multiple QTL models on the *M. guttatus* x *M. nudatus* linkage map. Points on each LOD interval indicate QTL peak positions.

The sum of these loci exceeded the parental divergence in node of first flower and first internode length (381% and 111%, respectively), and explained most of the parental divergence in corolla width (66%), corolla length (96%), leaf shape (88%), and a modest percentage of leaf area (24%). For traits where the parental lines did not significantly differ (flowering time and height), allelic effects were in the direction of species divergence for 5 out of 9 QTL.

### Loci contributing to traits associated with serpentine endemism and reproductive isolation overlapped

As predicted from the phenotypic correlations in the F2 hybrids (Figure 3), loci contributing to multiple traits had overlapping 1.5-LOD intervals in five genomic regions (Table 2, Figure 4). Four genomic regions had overlapping loci contributing to 2-3 traits, while one region had loci contributing to 5 traits. QTL overlap was often, but not always, between putatively adaptive and reproductive isolating traits. For example, QTL contributing to corolla width and leaf area overlapped on chromosome 5 (the corolla width QTL also overlaps with a height QTL, but the height QTL does not overlap with the leaf area QTL on chromosome 5; Table 2, Figure 4). QTL contributing to height and corolla width overlapped on one arm of chromosome 8, QTL contributing to height at flowering, leaf shape, and corolla width overlapped on a separate arm of chromosome 8, and QTL contributing to first internode length and node of first flower overlapped on chromosome 10 (Table 2, Figure 4). One region on chromosome 11 contained 5 overlapping loci primarily contributing reproductive isolation between species (life history and flower size differences, Figure 4).

### Allelic effects at overlapping QTL were mostly in the direction of species divergence

In the five genomic regions where QTL overlapped, QTL effects were largely in the direction of species divergence, suggesting that genetic correlations facilitated divergence in this species pair. In the overlapping region on chromosome 10, F2s homozygous for *M. guttatus* alleles flowered at later nodes and had shorter first internodes (Table 3). In the overlapping region on chromosome 5, F2s homozygous for *M. guttatus* alleles were taller, had wider flowers, and larger leaves (Table 3). Chromosome 8 had two overlapping regions on opposite ends of the chromosome. In the first overlapping region on chromosome 8, allelic effects differed relative to the direction as species divergence: F2s homozygous for *M. guttatus* alleles were shorter and had wider flowers (Table 3). In the second overlapping region on chromosome 8, allelic effects were all in the direction of species divergence: F2s homozygous for *M. guttatus* alleles were taller, had wider corollas and rounder leaves (Table 3). Finally, in the overlapping region on chromosome 11, F2s homozygous for *M. guttatus* alleles flowered later, flowered at later nodes, had shorter first internodes, wider and longer corollas (Table 3).

### Floral and leaf trait divergence is likely due to directional selection

We were only able to apply the QTL sign test with equal effects on traits where we identified more than six QTL, plant height at flowering and leaf shape. For plant height at flowering, allelic effects at 4 out of 8 QTL were in the direction of species divergence. For leaf shape, allelic effects at 3 out of 6 QTL were in the direction of species divergence. We could not reject the null hypothesis of neutral divergence between species for these two traits (*p* > 0.05), although this test is conservative (all six leaf shape QTL, and 7 out of 8 height QTL would need to be in the direction of species divergence to reject the null hypothesis).

We were not constrained by the number of QTL for the *v*-test and found support for a history of directional selection on the length of the first internode, corolla width, corolla length, leaf area, and leaf shape (Bonferroni adjusted *p* < 0.05; Table S1). We could not reject our null hypothesis of neutral divergence for days to flowering, node of first flower, and plant height at flowering (Bonferroni adjusted *p* > 0.05; Table S1).

### QTL contain homologs of genes regulating flowering time, flower size, and leaf traits

The 1.5 LOD regions for QTL we identified contain hundreds of annotated genes in the *M. guttatus* v2 reference genome (average: 403, range: 102-1065, Table S2). These regions contained homologs of *Arabidopsis thaliana* genes with known functions in regulating flowering time, growth, flower size, and leaf size and shape (Table S2). For example, the overlapping 1.5 LOD interval for flowering time, flowering node, and internode length on chromosome 11 includes a homolog of *CYCLIN-DEPENDENT KINASE G2* (Migut.K00703), a negative regulator of flowering that regulates alternative splicing of *FLOWERING LOCUS M* (Ma et al., 2015; Nibau et al. 2020). The overlapping 1.5 LOD interval for flowering node and internode length on chromosome 10 includes multiple homologs of flowering time regulators, including *APR6* (Migut.J01219; Deal et al., 2005), *MMP* (Migut.J01238; Golldack et al., 2002), *CTR1* (Migut.J00985; Achard et al., 2007), *SPY* (Migut.J01003; Jacobsen et al., 1996), *ABI5* (Migut.J01048; Wang et al., 2013), and *ADS1* (Migut.J01096; Sun et al., 2011) (Table S2).

Multiple candidate genes regulating organ growth and final organ size were located within the 1.5 LOD intervals of the flower size, leaf size and leaf shape QTL (Table S2). For example, the leaf shape QTL on chromosomes 6, 8, and 14 (QTL with effects in the direction of the parental divergence) contain homologs of genes influencing leaf shape (*ROTUNDIFOLIA LIKE 8* homolog Migut.F01456, Wen et al. 2004; *ROTUNDIFOLIA LIKE 3* homolog Migut.H02504, Kim et al. 1998) and size (*ARGOS* homolog Migut.N03014, Hu et al. 2003; *GIBBERELIN 20 OXIDASE 1* homolog Migut.F01163, Gonzalez et al. 2010) in *Arabidopsis*. The corolla width QTL on chromosomes 5, 8, and 11 include homologs of transcription factors involved in the control of petal size, including *BIGPETALp* (Migut.E00124; Szécsi et al., 2006), *AINTEGUMENTA* (Migut.H00323; Mizukami and Fischer, 2000), and *KLU* (Migut.K00730; Anastasiou et al., 2007). The overlapping QTL region contributing to both corolla width and leaf area on chromosome 5 includes pleiotropic candidate genes that affect petal size and leaf size or shape (*PAPS1* homolog Migut.E00384 [Vi et al. 2003] and *JAG* homolog Migut.E00347 [Ohno et al. 2004; Dinneny et al. 2004]). However, mutations at *PAPS1* that increase leaf size decrease flower size in *Arabidopsis*, and mutations at *JAG* that decrease petal size and cause leaf serration.

## Discussion

Edaphic variation generates strong divergent selection that contributes to local adaptation between soil ecotypes, and the evolution of new species. Edaphic variation may be more likely to contribute to the formation of new species, if traits associated with edaphic adaptation simultaneously contribute to reproductive isolation, or if these traits are genetically correlated through pleiotropy or linkage. Here we found that differences in putatively adaptive and reproductive isolating traits between a serpentine soil endemic, *M. nudatus*, and its closely related widespread congener, *M. guttatus*, are genetically correlated through linkage or pleiotropy (Figure 4). Our two effect size estimates, the percent of F2 trait variance explained and the percent of the parental trait divergence explained yield complementary information about the genetic architecture of adaptive divergence (Table 3). In terms of the percent variance explained, few large effect loci contributed to divergence in a subset of traits, but many small effect loci contributed to divergence in all traits. In terms of the percent of the parental divergence explained, few loci had intermediate to large effects on all traits, but parental divergence was not fully explained for most traits, suggesting that additional small effect loci contribute. Our analyses of selection supported the hypothesis that a history of directional selection likely contributed to species divergence in life history, floral, and leaf traits (Table S1). The distribution of effect sizes we observed may reflect the more gradual environmental variation experienced by populations after initial adaptation to serpentine soils. Alleles from serpentine endemic *M. nudatus* were often recessive, which could reflect adaptive loss of function alleles in *M. nudatus* (Monroe et al., 2021), or may be related to *M. nudatus’* demographic history, since speciation likely began with the initial colonization of small population on serpentine soils (Slatkin, 1996).

### Major and minor QTL contribute to traits associated with serpentine endemism

Many soil endemics, including *M. nudatus*, can grow on surrounding soils, suggesting that specialization is not due to unique abiotic requirements for growth (Krukeberg, 1954; Baskin & Baskin 1988). Thus, biotic factors, like competition and herbivory, are hypothesized to restrict their distributions. The repeated evolution of similar traits, like xeromorphic (small, narrow, or succulent) leaves and short statures, in the floras of unique soil types suggest that these traits adaptive (Krukeberg, 1954; Escudero et al., 2014), and may partially explain endemism, as smaller sizes and higher light requirements may reduce competitive abilities off serpentine soils (Krukeberg, 1954; Baskin & Baskin, 1988). Our analyses suggest that a history of consistent directional natural selection is likely responsible for species divergence in leaf area and shape, although we could not reject neutrality for plant height at flowering (Table S1). Nevertheless, *M. nudatus’* small size might also contribute to its endemism, especially if its small size reduces its competitive ability off serpentine soils. Further, the evolution of serpentine tolerance could directly contribute to endemism in *M. nudatus* if alleles that increase serpentine tolerance decrease competitive ability off or on less harsh serpentine soils. We identified eight small to large effect QTL contributing to plant height at flowering, the largest of which overlapped with a genomic region associated with serpentine tolerance between locally adapted populations of *M. guttatus* (Selby & Willis, 2018). However, allelic effects at this locus are in the opposite direction of species divergence; the *M. nudatus* homozygote at the chromosome 13 QTL is taller at flowering than the *M. guttatus* homozygote. Given that our hybrids were grown in individual pots without above or below ground competition, it is likely that loci influencing plant size will differ in natural environments. Furthermore, our parental lines did not significantly differ in height at flowering in our greenhouse common garden, suggesting that observed height differences between species in the field are environment dependent or that height differences are more pronounced later in the life cycle, which we did not measure. In addition, our parental lines were both from serpentine tolerant populations, and height differences are greater between *M. nudatus* and off serpentine populations of *M. guttatus* (Selby & Willis, 2018).

Serpentine soils are not only chemically challenging for plant species but are physically challenging due to their shallowness, rockiness and low water holding capacity. It is perhaps not surprising that plants that live on serpentine have repeatedly evolved xeromorphic foliage. Consistent with this pattern, *M. nudatus’* has very small narrow leaves, a unique trait in the *M. guttatus* species complex. We previously used a candidate QTL approach to identify genomic regions contributing to divergence in leaf shape between *M. nudatus* and *M. guttatus* (Ferris et al. 2015). In that study, we genotyped F2s derived from a cross between an allopatric non-serpentine *M. guttatus* population and *M. nudatus* at markers within three regions associated with leaf shape differences between a lobed leaf granite endemic *M. laciniatus* and allopatric round leaved *M. guttatus*. We also found significant associations with leaf shape at two markers on chromosomes 2 and 4. Only one of the six QTL, on chromosome 4, identified in our current study overlapped with a previously identified QTL. However allelic effects at this locus were in the opposite direction of the parental divergence – *M. nudatus* homozygotes had rounder leaves than *M. guttatus* homozygotes. In a subsequent QTL mapping study between *M. laciniatus* and a sympatric population of *M. guttatus*, different loci were identified contributing to leaf shape differences (Ferris et al. 2017). Two QTL from that subsequent study, on chromosomes 5 and 8, overlap with a leaf area and leaf shape QTL identified in our current study. Allelic effects at these loci were in the same direction as parental divergence – *M. nudatus* homozygotes had smaller and narrower leaves at the chromosome 5 and 8 QTL, respectively. Although these regions are associated with different leaf phenotypes (lobed vs small vs narrow leaves), QTL overlap suggests that these regions may contain genes involved in the repeated evolution of leaf size and shape in edaphic endemics.

While our QTL regions contain hundreds of annotated genes (Table S2), these regions contained promising homologs of *Arabidopsis thaliana* genes with known functions regulating leaf size and shape (Table S2). For example, our chromosome 5 leaf area QTL contains a homolog of *CYCLIN D1;1*, a cyclin dependent kinase that regulates leaf cell division and size (Cho et al., 2004; Zhang et al., 2019). Our chromosome 8 leaf shape QTL contains a homolog of *TCP3* (Migut.H01974), a transcription factor that regulates heterochronic leaf differentiation (Efroni et al., 2008), and a homolog of *ROT3*, a cytochrome P-450 enzyme that regulates polar elongation of leaf cells (Kim et al., 1998).

### Major and minor QTL contribute to prezygotic reproductive isolation

Although flowering time differences between *M. nudatus* and *M. guttatus* have been observed in the greenhouse (Sianta & Kay, 2021) and in the field (Selby et al., 2014), our parental lines did not significantly differ in our greenhouse common garden. Despite no significant difference in flowering time, we observed a significant difference in the developmental timing of allocation to reproductive growth between the *M. nudatus* and *M. guttatus* parental lines. *M. nudatus* transitioned to flowering at an earlier developmental stage, reflected by elongation of the first internode (bolting) and by the production of flowers at earlier nodes (Table 1). In our selection analysis, we found support for consistent directional natural selection on earlier bolting but could not reject the null hypothesis of neutrality for flowering time or the node of first flower. All three traits shared a major QTL on chromosome 11, suggesting that while interspecific flowering time differences are context dependent, shifts in developmental timing are genetically correlated with flowering time variation. *M. nudatus* has a shorter critical photoperiod requirement for flowering, compared to sympatric populations of *M. guttatus* (8 vs 13 hours of light per day required for flowering), but these species do not always differ in flowering time under constant long days (Freidman & Willis, 2013), possibly underlying the long co-flowering period in the field (Toll & Willis, 2018; Toll & Lowry, 2022).

Flower size differences between *M. nudatus* and *M. guttatus* contribute to floral mechanical isolation between these species (Gardner and MacNair 2000). *M. nudatus* is primarily pollinated by small-bodied bees, whereas *M. guttatus* is primarily pollinated by larger bodied bees, however pollinators frequently transition between species (Gardner & MacNair 2000; Koski et al., 2015; Toll, personal observation). Gardner & MacNair (2000) observed that small-bodied bees transitioning from *M. nudatus* to *M. guttatus* flowers were able to avoid touching *M. guttatus’* stigma while collecting pollen due to *M. guttatus’* large flower size. Our selection analysis supported the hypothesis that a history of consistent divergent natural selection contributed to species divergence in corolla width and length, although the source of this selection is not known. Divergence in flower size may have been driven by selection to reduce costly interspecific hybridization (Servedio & Noor, 2003), or as a drought adaptation to serpentine soils. Given that corolla width and length had one of the highest observed correlations in the F2s (Figure 3), and we only detected a single QTL contributing to corolla length with a similar LOD peak and identical LOD interval to a corolla width QTL, it is unlikely that directional selection contributed to divergence in these traits independently. However, we cannot distinguish which trait was the direct target of directional selection with this experiment, or whether correlations with a different unmeasured trait resulted in indirect selection on flower size.

We found that major and minor QTL contribute to divergence in flower size between *M. nudatus* and *M. guttatus*, consistent with genetic mapping studies across the *M. guttatus* species complex (Fishman et al., 2002; Ferris et al., 2017). The corolla width QTL we identified contain promising candidate genes (Table S2), including a homolog of *BIGPETALp*, a basic helix-loop-helix gene that controls petal size (Szécsi et al., 2006), a homolog of *AINTEGUMENTA* a transcription factor that regulates floral organ cell number and cell size (Mizukami & Fischer, 2000), and *KLU* a cytochrome P450 enzyme that promotes petal growth (Anastasiou et al., 2007). All the flower size QTL we detected co-localize with previously identified loci, suggesting that the same genes might be involved in intra- and inter-specific divergence in the *M. guttatus* species complex. The chromosome 5 QTL we identified co-localizes with an interspecific flower size QTL between *Mimulus laciniatus* and *M. guttatus* (Ferris et al. 2017) and *Mimulus nasutus* and *M. guttatus* (Fishman et al., 2002). Both chromosome 8 QTL we identified overlap with flower size QTL between annual and perennial ecotypes of *M. guttatus* (Hall et al., 2006) and between *M. laciniatus* and *M. guttatus* (Ferris et al., 2017). Finally, the chromosome 11 QTL we identified co-localizes with flower size QTL between *M. nasutus* and *M. guttatus* (Fishman et al., 2002).

### Genetic correlations may have facilitated adaptation and speciation

Soils that are chemically and physically unusual, like serpentine and gypsum, cover a low percentage of the earth’s surface but harbor a high percentage of endemic species (Brady et al., 2005; Escudero et al., 2014). Despite the high diversity and endemism on serpentine soils, tolerator species (species with populations on and off serpentine soils) outnumber endemics four to one (Anacker, 2014). One reason why serpentine endemics may be less common than tolerator species is because strong divergent selection across soil boundaries can readily generate genetic divergence between populations but is unlikely to result in the evolution of reproductive isolation on its own (Bolnick & Fitzpatrick, 2007). A period of allopatry, where small populations are genetically isolated on edaphic islands, is likely necessary to complete the process of speciation. Genetic correlations between traits that increase assortative mating and fitness under divergent selection across soil boundaries may facilitate the evolution of adaptation and reproductive isolation between ecotypes (Felsenstein, 1981; Hawthorne & Via, 2001). Following a period of allopatry, genetic correlations between adaptive and reproductive isolating traits also increase the probability of coexistence by maintaining species boundaries upon secondary contact (Rundle & Nosil, 2005).

*M. nudatus* and *M. guttatus* are reproductively isolated through several prezygotic isolating barriers, including microhabitat segregation (Toll & Willis, 2018; Toll et al., 2021; Toll & Lowry, 2022), flowering time differences (Sianta & Kay, 2021), flower size and pollinator differences (Gardner & MacNair, 2000), and a postzygotic isolating barrier, hybrid seed inviability (Gardner & MacNair, 2000, Oneal et al., 2016). We found that traits contributing to assortative mating and traits associated with the evolution of serpentine endemism are genetically correlated and may have facilitated divergence and/or coexistence in this species pair. Although our QTL mapping experiment cannot distinguish between pleiotropy and linkage, genomic regions with overlapping QTL often had similar peak positions and LOD profiles (Table 2, Figure 4). However, most of the candidate genes we identified in the overlapping 1.5-LOD regions on chromosomes 5, 8 and 11 have no published pleiotropic effects on flowering time, leaf, and floral traits. Thus, QTL overlap may be due to linkage among genes influencing these traits individually.

### M. nudatus alleles were more likely to be recessive

For most partially dominant and dominant QTL, the *M. nudatus* allele was recessive (14 out of 17, Table 3). This pattern contrasts with genetic mapping studies between predominantly self-fertilizing and outcrossing monkeyflower species, where no directional dominance was observed, and is unexpected because these species have similar mixed mating systems (Fishman et al., 2002; Ferris & Willis, 2017). The preponderance of recessive alleles in *M. nudatus* could be related to the types of substitutions fixed during divergence or could be related to its demographic history. Loss of function alleles are likely to be recessive and are consistent with the direction of phenotypic differences between species, as many studies describe gene knockouts with smaller organ sizes and dwarf phenotypes (Table S2). Although we do not know what the causal genes are in our QTL, much less the causal mutations, a promising direction for future studies is investigating whether candidate genes have obvious knockouts, including premature stops, radical AA substitutions, and indels in coding regions.

A non-mutually exclusive hypothesis is that founder events in *M. nudatus’* demographic history resulted in the increased fixation of beneficial recessive alleles. In outcrossing populations, adaptation from new mutations is biased against the fixation of recessive alleles (Haldane, 1927), while adaptation from standing variation is unbiased with respect to dominance (except for fully dominant loci, Orr & Betancourt, 2001). The probability of fixation of beneficial recessive alleles increases with self-fertilization (Charlesworth, 1992), but *M. nudatus* and *M. guttatus* both have similar mixed mating systems (Toll et al., 2021). For a given initial allele frequency, the probability of fixation of a beneficial recessive allele decreases with decreasing population size (Kimura 1962). However, fixation probabilities increase with initial allele frequencies and allele frequencies are initially higher in small populations provided those alleles are not eliminated by drift. When followed by a period of rapid growth, those recessive alleles are at higher frequency than they would have been if population sizes remained constant, increasing their probability of fixation (Slatkin, 1999). This demographic scenario is likely during the evolution of edaphic endemics, where small populations initially colonize new edaphic environments and followed by a period of rapid population growth (Kruckeberg & Rabinowitz, 1982). Future population genetic studies in *Mimulus nudatus* may clarify whether this scenario is likely and can explain the patterns of dominance we observed.

### Conclusion

Physically and chemically unique soils harbor floras characterized by narrowly endemic species. The evolution of these edaphic endemics is influenced by the genetic architecture of divergence across soil boundaries. Correlations between traits that increase edaphic adaptation and assortative mating can increase the probability of speciation and coexistence upon secondary contact. Consistent with theory about the effect size distribution of adaptive substitutions, we found that few large effect and many small effect loci contribute to divergence between the serpentine endemic, *M. nudatus*, and co-occurring serpentine adapted populations of its widespread close relative *M. guttatus*. Selection analyses support the hypothesis that consistent directional natural selection contributed to species divergence in the developmental timing of bolting, flower size, and leaf size and shape. For partially dominant and dominant QTL, alleles from the serpentine endemic were mostly recessive, potentially due to the types of substitutions fixed during divergence or founder events in its demographic history. We also found that QTL contribute to traits associated with serpentine endemism (small leaves and stature) and reproductive isolation (flower size and flowering time) overlap, either due to pleiotropy or linkage. These overlapping QTL contain many genes and thus, future fine mapping studies are necessary to distinguish between pleiotropy and linkage. Overall, our study suggests that genetic correlations facilitated the evolution of adaptation and speciation in this pair of species.

## Acknowledgments

The authors wish to thank Laryssa Baldridge, Jennifer Coughlan, Kyle Christie, Meng Chen, Kathleen Donohue, Katie Ferris, Annie Jeong, Eric LoPresti, David Lowry, Paul Magwene, Tom Mitchell-Olds, Elen Oneal, Jessica Selby, Mark Rausher, Ashley Troth, and Carrie Wessinger for advice and feedback throughout this project. We thank Natalie Knox and Brian Weill for assisting with plant care in the greenhouse. This work was supported by National Science Foundation (NSF) Grants #1354688 and #1558113 awarded to J.H. Willis and an NSF Doctoral Dissertation Improvement Grant #1406952 awarded to K. Toll and J. H. Willis.

## Data Accessibility

Raw sequence reads will be deposited in the SRA (BioProject XXX). Individual genotype data, phenotype data, and R code will be available on Dryad upon acceptance (Toll & Willis 2022).

## Author contributions

KT and JHW designed the QTL mapping experiment, KT performed the experiment, analyzed the data, and wrote the manuscript.

