## SupplementalMaterials for "The genetic basis of traits associated with the evolution of serpentine endemism in monkeyflowers"

**Supplemental materials**

**Figure S1.** F2 genotype frequencies and regions with significant segregation distortion in the F2s.

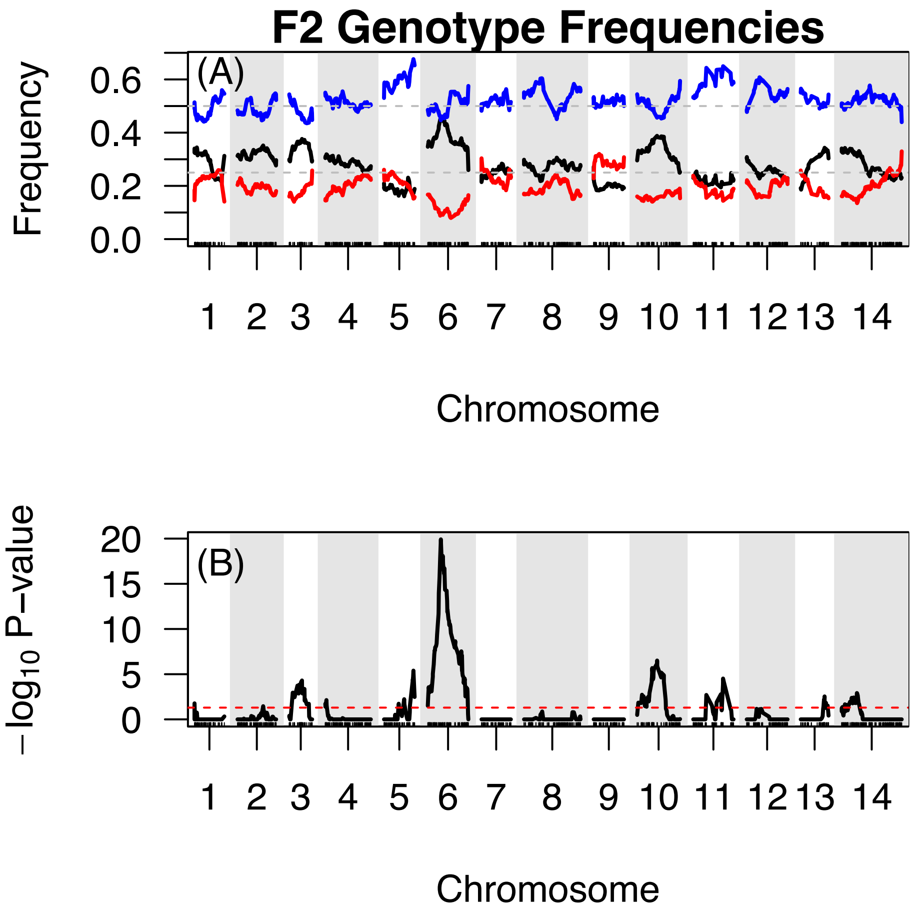


**Table S1.** *v*-test of consistent directional selection with a genetic cross (Fraser, 2020).

| Trait | Variance Between Parents | F2 Variance | *v* | *p*-value | Bonferonni adjusted *p*-value |
|---|---|---|---|---|---|
| Days to Flowering | 13.383 | 23.03508 | 0.6809093 | 0.4092738 | 1 |
| Node of First Flower | 2.38235 | 1.051463 | 1.5161 | 0.2182106 | 1 |
| First Internode Length | 119.335 | 3.609157 | 21.28068 | 3.97E-06 | **3.18E-05** |
| Height | 7.296 | 13.66345 | 0.3794183 | 0.5379147 | 1 |
| Corolla Width | 7.296 | 0.08309566 | 74.19306 | 7.08E-18 | **5.66E-17** |
| Corolla Length | 1.21637 | 0.1542013 | 8.095373 | 0.00443784 | **3.55E-02** |
| Leaf Area | 354.11 | 4.627331 | 168.1571 | 1.87E-38 | **1.50E-37** |
| Leaf Shape | 0.4012 | 0.01314662 | 20.98773 | 4.62E-06 | **3.70E-05** |

**Table S2.** Candidate genes in QTL regions. QTL direction refers to the direction of the QTL effect relative to species divergence (+ is the same direction as species divergence, - is the opposite direction relative to species divergence). Gene annotations are from the *M. guttatus* v2 reference genome (Hellsten et al. 2013), gene homolog information is from The *Arabidopsis* Information Resource (Berardini et al., 2015).

| **Trait** | **QTL direction** | **Chromosome** | **Annotated gene in 1.5-LOD interval** | ***M. guttatus* locus name** | ***A. thaliana* locus name** | ***A. thaliana* other name** | **Citation** |
|---|---|---|---|---|---|---|---|
| **Days to Flowering** | + | 11 | Migut.K00439-Migut.K00707 | Migut.K00470 | AT1G04870 | PROTEIN ARGININE METHYLTRANSFERASE 10 (ATPRMT10) | Niu et al. 2007 |
| | | | | Migut.K00558 | AT4G29230 | NAC DOMAIN CONTAINING PROTEIN 75 (ANAC075) | Fujiwara et al. 2016 |
| | | | | Migut.K00703 | AT1G67580 | CYCLIN-DEPENDENT KINASE G2 (CDKG2) | Ma et al. 2015; Nibau et al. 2020 |
| **Node of First Flower** | + | 10 | Migut.J00954-Migut.J01278 | Migut.J00985 | AT5G03730 | CONSTITUTIVE TRIPLE RESPONSE 1 (ATCTR1) | Achard et al. 2007 |
| | | | | Migut.J01003 | AT3G11540 | SPINDLY (SPY) | Jacobson et al. 1996 |
| | | | | Migut.J01048 | AT2G36270 | ABA INSENSITIVE 5 (ABI5) | Wang et al. 2013 |
| | | | | Migut.J01096 | AT4G29140 | ACTIVATED DISEASE SUSCEPTIBILITY 1 (ADS1) | Sun et al 2011 |
| | | | | Migut.J01219 | AT3G33520 | ACTIN-RELATED PROTEIN6 (ARP6) | Deal et al. 2005 |
| | | | | Migut.J01238 | AT1G70170 | MATRIX METALLOPROTEINASE (AT2-MMP) | Golldack et al. 2002 |
| | + | 11 | Migut.K00640-Migut.K00741 | Migut.K00703 | AT1G67580 | CYCLIN-DEPENDENT KINASE G2 (CDKG2) | Ma et al. 2015; Nibau et al. 2020 |
| **First Internode Length** | + | 10 | Migut.J01038-Migut.J01308 | same as node of first flower | | | |

| | | | | | | | |
|---|---|---|---|---|---|---|---|
| | + | 11 | Migut.K00640-Migut.K00741 | same as node of first flower | | | |
| **Height** | - | 2 | Migut.B00933-Migut.B01168 | Migut.B01040 | AT3G07610 | INCREASE IN BONSAI METHYLATION 1 (IBM1) | Saze et al. 2008 |
| | | | | Migut.B01057 | AT3G07610 | INCREASE IN BONSAI METHYLATION 1 (IBM1) | Saze et al. 2008 |
| | + | 5 | Migut.E00468-Migut.E01348 | Migut.E00579 | AT4G21200 | ARABIDOPSIS THALIANA GIBBERELLIN 2-OXIDASE 8 (ATGA2OX8) | Schomburg et al. 2003 |
| | | | | Migut.E00766 | AT1G79460 | GA REQUIRING 2 (GA2); ARABIDOPSIS THALIANA ENT-KAURENE SYNTHASE (ATKS) | Yamaguchi et al. 1998 |
| | | | | Migut.E00897 | AT1G79460 | GA REQUIRING 2 (GA2); ARABIDOPSIS THALIANA ENT-KAURENE SYNTHASE (ATKS) | Yamaguchi et al. 1998 |
| | | | | Migut.E01186 | AT1G79460 | GA REQUIRING 2 (GA2); ARABIDOPSIS THALIANA ENT-KAURENE SYNTHASE (ATKS) | Yamaguchi et al. 1998 |
| | | | | Migut.E01185 | AT4G02780 | GA REQUIRING 1 (GA1); | Koorneef and van der Veen 1980 |
| | - | 8a | Migut.H00001-Migut.H00739 | Migut.H00183 | AT5G66730 | ENHYDROUS (ENY); IDD1 (INDETERMINATE DOMAIN 1) | Fukazawa et al. 2014 |
| | | | | Migut.H00297 | AT5G65640 | NO FLOWERING IN SHORT DAY (NFL) | Sharma et al 2016 |
| | | | | Migut.H00468 | AT5G66880 | SUCROSE NONFERMENTING 1 (SNF1)-RELATED PROTEIN KINASE 2-3 (SNRK2-3) | Fujii and Zhu 2009 |
| | | | | Migut.H00592 | AT1G50420 | SCARECROW-LIKE 3 (SCL3) | Zhang et al. 2011 |
| | | | | Migut.H00528 | AT1G67260 | TCP DOMAIN PROTEIN 1 (TCP1) | Koyama et al 2010 |
| | | | | Migut.H00683 | AT5G51810 | GIBBERELLIN 20 OXIDASE 2 (GA20OX2) | Rieu et al. 2008 |
| | | | | Migut.E00409 | AT1G71090 | PIN-LIKES 2 (PILS2) | Barbez et al. 2012 |

| | | | | | | | |
|---|---|---|---|---|---|---|---|
| | | | | Migut.H00419 | AT5G13680 | ABA-OVERLY SENSITIVE 1 (ABO1) | Nelissen et al 2005 |
| | | | | Migut.H00583 | AT3G02260 | CORYMBOSA1 (CRM1) | Yamaguchi et al. 2007 |
| | | | | Migut.H00584 | AT3G02260 | CORYMBOSA1 (CRM1) | Yamaguchi et al. 2007 |
| | | | | Migut.H00589 | AT5G60910 | AGAMOUS-LIKE 8 (AGL8); FRUITFULL (FUL) | Bemer et al. 2017 |
| | | | | Migut.H00694 | AT5G63980 | FIERY1 (FRY1) | Kim and Arnim 2009 |
| | - | 8b | Migut.H01279-Migut.H02130 | Migut.H01362 | AT1G30040 | GIBBERELLIN 2-OXIDASE 2 (ATGA2OX2, GA2OX2) | Wang and Li 2006 |
| | | | | Migut.H01424 | AT4G18710 | BRASSINOSTEROID-INSENSITIVE 2 (BIN2) | Li et al. 2001 |
| | | | | Migut.H01518 | AT3G62980 | TRANSPORT INHIBITOR RESPONSE 1 (ATTIR1, TIR1) | Ruegger et al. 1998 |
| | | | | Migut.H01552 | AT4G21200 | GIBBERELLIN 2-OXIDASE 8 (GA2OX8) | Schomburg et al. 2003 |
| | | | | Migut.H01566 | AT5G05690 | CONSTITUTIVE PHOTOMORPHOGENIC DWARF (CPD) | Tanaka et al. 2005 |
| | | | | Migut.H01666 | AT1G14920 | GIBBERELLIC ACID INSENSITIVE (GAI) | Peng et al 1999 |
| | | | | Migut.M01872 | AT2G34650 | ABRUPTUS (ABR); PINOID (PID) | Christensen et al. 2000 |
| | | | | Migut.H01910 | AT5G54380 | THESEUS1 (THE1) | Guo et al. 2009 |
| | | | | Migut.H02079 | AT1G30040 | GIBBERELLIN 2-OXIDASE 2 (ATGA2OX2, GA2OX2) | Wang and Li 2006 |
| | + | 9 | Migut.I00683-Migut.I00994 | Migut.I00703 | AT2G20000 | HOBBIT (HBT) | Perez-Perez et al. 2007 |
| | | | | Migut.I00709 | AT1G80680 | SUPPRESSOR OF AUXIN RESISTANCE 3 (SAR3) | Parry et al. 2006 |
| | | | | Migut.I00761 | AT3G15880 | TOPLESS-RELATED 4 (TPR4) | Garner et al 2021 |

| | | | | | | | |
|---|---|---|---|---|---|---|---|
| | | | | Migut.I00834 | AT1G15550 | ARABIDOPSIS THALIANA GIBBERELLIN 3 BETA-HYDROXYLASE 1 (ATGA3OX); GA REQUIRING 4 (GA4) | Mitchum et al. 2006 |
| | + | 11 | Migut.K00001-Migut.K00300 | Migut.K00023 | AT2G41940 | ZINC FINGER PROTEIN 8 (ZFP8) | Joseph et al 2014 |
| | | | | Migut.K00202 | AT4G18710 | BRASSINOSTEROID-INSENSITIVE 2 (BIN2) | Li et al. 2001 |
| | | | | Migut.K00276 | AT4G30410 | IBH1-LIKE 1 (IBL1) | Zhipanova et al. 2014 |
| | - | 13 | Migut.M00868-Migut.M01932 | Migut.M00997 | AT1G78700 | BES1/BZR1 HOMOLOG 4 (BEH4) | Wang et al. 2002 |
| | | | | Migut.M01017 | AT4G18710 | BRASSINOSTEROID-INSENSITIVE 2 (BIN2) | Li et al. 2001 |
| | | | | Migut.M01216 | AT3G62980 | TRANSPORT INHIBITOR RESPONSE 1 (ATTIR1, TIR1) | Ruegger et al. 1998 |
| | | | | Migut.M01278 | AT4G21200 | GIBBERELLIN 2-OXIDASE 8 (GA2OX8) | Schomburg et al. 2003 |
| | | | | Migut.M01279 | AT4G21200 | GIBBERELLIN 2-OXIDASE 8 (GA2OX8) | Schomburg et al. 2003 |
| | | | | Migut.M01506 | AT1G79460 | GA REQUIRING 2 (GA2); ARABIDOPSIS THALIANA ENT-KAURENE SYNTHASE (ATKS) | Yamaguchi et al. 1998 |
| | | | | Migut.M01507 | AT1G79460 | GA REQUIRING 2 (GA2); ARABIDOPSIS THALIANA ENT-KAURENE SYNTHASE (ATKS) | Yamaguchi et al. 1998 |
| | | | | Migut.M01835 | AT5G54510 | DWARF IN LIGHT 1 (DFL1); GRETCHEN HAGEN3.6 (GH3.6) | Nakazawa et al. 2008 |
| | | | | Migut.M01862 | AT1G30330 | AUXIN RESPONSE FACTOR 6 (ARF6) | Nagpal et al. 2005 |
| | | | | Migut.M01872 | AT2G34650 | ABRUPTUS (ABR); PINOID (PID) | Christensen et al. 200 |
| | | | | Migut.M01900 | AT1G30570 | HERCULES RECEPTOR KINASE 2 (HERK2) | Guo et al. 2009 |
| | + | 14 | Migut.N00993-Migut.N01143 | Migut.N01009 | AT4G33430 | BRASSINOSTEROID-INSENSITIVE 2 (BIN2) | Li et al. 2001 |

| | | | | | | | |
|---|---|---|---|---|---|---|---|
| | | | | Migut.N01109 | AT3G03730 | KINK SUPPRESSED IN BZR1-1D 4 (KIB4) | Zhu et al. 2017 |
| | | | | Migut.N01118 | AT5G65810 | COTTON GOLGI-RELATED 3 (CGR3) | Kim et al. 2005 |
| **Corolla Width** | + | 5 | Migut.E00001-Migut.E00560 | Migut.E00124 | AT1G59640 | BIG PETAL P (BPEp) | Szecsi et al. 2006 |
| | | | | Migut.E00302 | AT4G24150 | GROWTH-REGULATING FACTOR 8 (ATGRF8, GRF8) | Krizek and Anderson 2013 |
| | | | | Migut.E00347 | AT1G68480 | JAGGED (JAG) | Ohno et al. 2004; Dinneny et al. 2004 |
| | | | | Migut.E00384 | AT1G17980 | POLY(A) POLYMERASE 1 (PAPS1) | Vi et al. 2013 |
| | + | 8a | Migut.H00171-Migut.H00516 | Migut.H00323 | AT4G37750 | AINTEGUMENTA (ANT) | Mizukami and Fischer 2000 |
| | | | | Migut.H00324 | AT2G22840 | GROWTH-REGULATING FACTOR 1 (ATGRF1, GRF1) | Krizek and Anderson 2013 |
| | | | | Migut.H00399 | AT5G67260 | CYCLIN D3;2 (CYCD3;2) | Dewitte et al. 2007 |
| | + | 8b | Migut.H01957-Migut.H02204 | Migut.H02192 | AT3G51390 | PROTEIN S-ACYL TRANSFERASE 10 (ATPAT10, PAT10) | Zhou et al. 2013 |
| | + | 11 | Migut.K00640-Migut.K00954 | Migut.K00730 | AT1G13710 | KLUH (KLU/CYP78A5) | Anastasiou et al 2007 |
| **Corolla Length** | + | 11 | Migut.K00640-Migut.K00954 | Migut.K00730 | AT1G13710 | KLUH (KLU/CYP78A5) | Anastasiou et al 2007 |
| **Leaf Area** | + | 5 | Migut.E00129-Migut.E00457 | Migut.E00135 | AT1G70170 | MATRIX METALLOPROTEINASE (MMP) | Golldack et al. 2002 |
| | | | | Migut.E00142 | AT1G70210 | CYCLIN D1;1 (CYCD1;1) | Cho et al. 2004; Zhang et al. 2018 |
| | | | | Migut.E00302 | AT4G24150 | GROWTH-REGULATING FACTOR 8 (ATGRF8, GRF8) | Kim et al. 2003 |
| | | | | Migut.E00336 | AT4G27950 | CYTOKININ RESPONSE FACTOR 4 (CRF4) | Raines et al. 2015 |

| | | | | | | | |
|---|---|---|---|---|---|---|---|
| | | | | Migut.E00376 | AT1G49540 | ELONGATOR SUBUNIT 2 (ELP2) | Nelissen et al. 2005 |
| | | | | Migut.E00384 | AT1G17980 | POLY(A) POLYMERASE 1 (PAPS1) | Vi et al. 2013 |
| | - | 6 | Migut.F01877-Migut.F02142 | Migut.F02013 | AT4G10090 | ELONGATOR SUBUNIT 6 (ELP6) | Zhou et al. 2009 |
| | | | | Migut.F02083 | AT1G79620 | VASCULAR-RELATED RLK 1 (VRLK1) | Huang et al 2018 |
| **Leaf Shape** | - | 4 | Migut.D01779-Migut.D01985 | Migut.D01823 | AT4G18750 | DEFECTIVELY ORGANIZED TRIBUTARIES 4 (DOT4) | Petricka et al. 2008 |
| | + | 6 | Migut.F01047-Migut.F01528 | Migut.F01163 | AT4G25420 | GIBBERELLIN 20 OXIDASE 1 (GA20OX1); GA REQUIRING 5 (GA5) | Gonzalez et al. 2010 |
| | | | | Migut.F01420 | AT5G05780 | RP NON-ATPASE SUBUNIT 8A (RPN8A) | Huang et al. 2006 |
| | | | | Migut.F01456 | AT2G39705 | DEVIL 11 (DVL11); ROTUNDIFOLIA LIKE 8 (RTFL8) | Narita et al. 2004; Wen et al 2004 |
| | + | 8 | Migut.H01654-Migut.H02559 | Migut.H01895 | AT3G20630 | TARANI (TNI, TTN6) | Karidas et al. 2015 |
| | | | | Migut.H01966 | AT1G53190 | TIE1-ASSOCIATED RING-TYPE E3 LIGASE1 (TEAR1) | Zhang et al. 2007 |
| | | | | Migut.H01974 | AT1G53230 | TEOSINTE BRANCHED 1, cycloidea and PCF transcription factor 3 (TCP3) | Efroni et al 2008; Koyama et al. 2017 |
| | | | | Migut.H02162 | AT4G38630 | REGULATORY PARTICLE NON-ATPASE 10 (RPN10) | Smalle et al. 2003 |
| | | | | Migut.H02192 | AT3G51390 | PROTEIN S-ACYL TRANSFERASE 10 (ATPAT10, PAT10) | Zhou et al. 2013 |
| | | | | Migut.H02340 | AT4G33950 | UCROSE NONFERMENTING 1-RELATED PROTEIN KINASE 2-6 (SNRK2.6) | Zheng et al. 2010 |
| | | | | Migut.H02355 | AT5G10270 | CYCLIN-DEPENDENT KINASE C;1 (CDKC;1) | cyclin-dependent kinase C;1 |

| | | | | | | | |
|---|---|---|---|---|---|---|---|
| | | | | Migut.H02359 | AT5G10250 | DEFECTIVELY ORGANIZED TRIBUTARIES 3 (DOT3) | Petricka et al. 2008 |
| | | | | Migut.H02431 | AT5G09970 | CYTOCHROME P450, FAMILY 78, SUBFAMILY A, POLYPEPTIDE 7 (CYP78A7) | Nobusawa et al. 2021 |
| | | | | Migut.H02504 | AT4G36380 | ROTUNDIFOLIA 3 (ROT3) | Kim et al. 1998 |
| | - | 9 | Migut.I00220-Migut.I00659 | Migut.I00266 | AT3G48430 | RELATIVE OF EARLY FLOWERING 6 (REF6) | Yu et al. 2008 |
| | | | | Migut.I00277 | AT5G50320 | ELONGATOR PROTEIN 3 (ELP3) | Kojima et al. 2011 |
| | | | | Migut.I00299 | AT1G73590 | PIN-FORMED 1 (PIN1) | Hay et al. 2006 |
| | | | | Migut.I00387 | AT5G06110 | ZUOTIN-RELATED FACTOR 1B (ZRF1B) | Feng et al. 2016 |
| | | | | Migut.I00474 | AT5G50320 | ELONGATOR PROTEIN 3 (ELP3) | Kojima et al. 2011 |
| | | | | Migut.I00499 | AT5G14170 | SWP73B | Sacharowski et al. 2015 |
| | | | | Migut.I00510 | AT1G53190 | TIE1-ASSOCIATED RING-TYPE E3 LIGASE1 (TEAR1) | Zhang et al. 2007 |
| | | | | Migut.I00526 | AT1G53190 | TIE1-ASSOCIATED RING-TYPE E3 LIGASE1 (TEAR1) | Zhang et al. 2007 |
| | | | | Migut.I00527 | AT1G53190 | TIE1-ASSOCIATED RING-TYPE E3 LIGASE1 (TEAR1) | Zhang et al. 2007 |
| | | | | Migut.I00528 | AT1G53190 | TIE1-ASSOCIATED RING-TYPE E3 LIGASE1 (TEAR1) | Zhang et al. 2007 |
| | | | | Migut.I00530 | AT1G53190 | TIE1-ASSOCIATED RING-TYPE E3 LIGASE1 (TEAR1) | Zhang et al. 2007 |
| | | | | Migut.I00613 | AT5G62000 | AUXIN RESPONSE FACTOR 2 (ARF2) | Okushima et al. 2005 |
| | | | | Migut.I00616 | AT5G13300 | SCARFACE (SFC) | Deyholos et al. 2000 |
| | - | 12 | Migut.L00954-Migut.L01141 | Migut.L00956 | AT1G17980 | POLY(A) POLYMERASE 1 (PAPS1) | Vi et al. 2013 |
| | | | | Migut.L01010 | AT1G17980 | POLY(A) POLYMERASE 1 (PAPS1) | Vi et al. 2013 |

| | | | | | | | |
|---|---|---|---|---|---|---|---|
| | | | | Migut.L01011 | AT1G17980 | POLY(A) POLYMERASE 1 (PAPS1) | Vi et al. 2013 |
| | | | | Migut.L01093 | AT1G13290 | DEFECTIVELY ORGANIZED TRIBUTARIES 5 (DOT5) | C2H2-like zinc finger protein |
| | | | | Migut.L01097 | AT1G68825 | DEVIL 5 (DVL5); ROTUNDIFOLIA LIKE 15 (RTFL15) | Wen et al 2004; Narita et al. 2004 |
| | | | | Migut.L01141 | AT1G17110 | UBIQUITIN-SPECIFIC PROTEASE 15 (UBP15) | Liu et al. 2008 |
| | + | 14 | Migut.N02753-Migut.N03038 | Migut.N03014 | AT3G59900 | AUXIN-REGULATED GENE INVOLVED IN ORGAN SIZE (ARGOS) | Hu et al. 2003 |
| | | | | Migut.N02798 | AT4G37750 | AINTEGUMENTA (ANT) | Mizukami and Fischer 2000 |
| | | | | Migut.N02791 | AT4G07410 | POPCORN (PCN) | Xiang et al. 2011 |
| | | | | Migut.N02803 | AT3G49720 | COTTON GOLGI-RELATED 3 (CGR2) | Weraduwage et al. 2016 |
| | | | | Migut.N02870 | AT4G24560 | UBIQUITIN-SPECIFIC PROTEASE 16 (UBP16) | Liu et al. 2008 |

12
